# Repolarisation-relaxation dyscoupling and TRPA1 activation permit systolic mechano-arrhythmogenesis

**DOI:** 10.1101/2024.02.21.581489

**Authors:** Breanne A. Cameron, Matthew R. Stoyek, Joachim Greiner, Rémi Peyronnet, Peter Kohl, T Alexander Quinn

**Author notes:** **CORRESPONDING AUTHOR** Breanne A. Cameron Institute for Experimental Cardiovascular Medicine Universitätsklinikum Freiburg, Universitäts-Herzzentrum Elsässer Straße 2Q, 79110 Freiburg, Germany.

## Abstract

**Background:** The heart’s mechanical state feeds back to its electrical activity, potentially contributing to arrhythmias (mechano-arrhythmogenesis, MAR). MAR has been mechanistically explained during electrical diastole, when cardiomyocytes are at their resting membrane potential. Conversely, during electrical systole, cardiomyocytes appear to be protected from MAR, even as membrane potential and cytosolic calcium concentration ([Ca^2+^]_i_) are simultaneously restored to resting levels during repolarisation (repolarisation-relaxation coupling, RRC). Yet, systolic MAR has been reported in ischaemic myocardium, with unclear underlying mechanisms.

**Methods:** Rabbit left ventricular cardiomyocytes were electrically paced and exposed to a simulated ischaemia solution (including hyperkalaemia, acidosis, and block of oxidative phosphorylation) or pinacidil (to simulate ischaemia-induced opening of ATP-sensitive potassium [K_ATP_] channels), with or without glibenclamide (to block K_ATP_ channels). RRC was assessed by simultaneous measurement of membrane voltage and [Ca^2+^]_i_ dynamics with fluorescence imaging. Acute stretch at increasing magnitudes was applied using carbon fibres, with stretch timed to diastole or late systole. Stretch mechanics and the incidence of MAR was assessed by video-based measurement of sarcomere length. Mechanisms contributing to MAR were assessed by buffering [Ca^2+^]_i_ (BAPTA-AM), stabilising ryanodine receptors (dantrolene), non-specifically blocking mechano-sensitive channels (streptomycin), activating (AITC) or blocking (HC-030031) transient receptor potential kinase ankyrin 1 channels (TRPA1), or chelating (NAC) or blocking production of (DPI) reactive oxygen species (ROS).

**Results:** It was reconfirmed that MAR during physiological RRC is rare, while ischaemia- or pharmacologically-induced RRC dyscoupling generates a vulnerable period for systolic MAR. This systolic MAR depends on TRPA1, [Ca^2+^]_i_, and ROS, which contribute to stretch-induced excitation and arrhythmia sustenance. An increase in systolic MAR can be prevented by mitigating RRC dyscoupling with K_ATP_ channel block, or by blocking TRPA1, buffering [Ca^2+^]_i_, or reducing ROS.

**Conclusion:** RRC dyscoupling may be arrhythmogenic in ischaemia and other pathologies associated with systolic MAR, and TRPA1 may be a novel anti-arrhythmic target.

## INTRODUCTION

Feedback is an essential element of biological systems’ function and fundamental to the adaptation of physiological activity to varying demands. A prime example is found in the heart, an electro-mechanical pump whose electrical excitation causes mechanical contraction of cardiomyocytes (CM), involving a feedforward mechanism of electrically triggered calcium (Ca^2+^) release from intracellular stores (excitation-contraction coupling, ECC) and mechano-sensitive feedback mechanisms by which the mechanical state of CM affects their electrical activity (mechano-electric coupling, MEC).^1^ Although MEC is important for maintaining and fine-tuning normal cardiac function,^2^ it can also contribute to mechano-arrhythmogenesis (MAR)^3^ by causing aberrant electrophysiological behaviour, in particular during cardiac diseases that are associated with altered homogeneity of passive and active mechanics.^4^ While the influence of MEC on the heart’s electrical activity is well established,^1^ the mechanisms and molecular identity of specific driver(s) underlying MAR are unclear.^5^

MEC-driven excitation has been reliably explained during ‘electrical diastole’ (here used to refer to the phase of the cardiac cycle in which CM are at their resting membrane potential, V_m_). This has been shown to involve mechano-sensitive ion channels, whose activation depolarises CM, and mechanical modulation of intracellular Ca^2+^ handling,^1^ such as a stretch-induced increase in sarcoplasmic reticulum Ca^2+^ release^6^ or stretch-and-release related reductions in myofilament Ca^2+^binding.^7^ On the other hand, under physiological conditions CM appear to be well-protected from MEC-driven arrhythmogenesis during ‘electrical systole’ (*i.e.,* while the action potential [AP] is occurring). This includes both early (*i.e.*, ECC and plateau phases) and late electrical systole, the period during which V_m_ and free cytosolic Ca^2+^ concentration ([Ca^2+^]_i_) are concurrently being restored to resting values. This restoration is achieved by progressive V_m_ repolarisation (which prevents further Ca^2+^ influx through voltage-gated channels, while simultaneously increasing CM excitability) and reduction in [Ca^2+^]_i_ (*via* sequestration back into intracellular stores and extrusion to the cell exterior), leading to mechanical relaxation. This is a well-coordinated process, henceforth referred to as ‘repolarisation-relaxation coupling’ (RRC). Protection from systolic MAR has been attributed to transient electrical refractoriness and to the tight coordination of electro-mechanical activation and recovery in CM. In pathological states, however, a dissociation of electro-mechanical dynamics may weaken this protection.^4^

One such pathology is ischaemia, which causes dyscoupling of the normally well-synchronised late-systolic recovery of V_m_ and [Ca^2+^]_i_ during RRC. This dyscoupling can occur through activation of ATP-sensitive potassium (K_ATP_) channels with reduced oxygen availability. K_ATP_ channels conduct an outward current over the entire working range of CM V_m_, resulting in early repolarisation (*i.e.,* a shortened AP) that is not matched by an equally pronounced shortening of the [Ca^2+^]_i_ transient. This causes a corresponding increase in the duration of RRC (T_RRC_, the time between the recovery of CM excitability and return of [Ca^2+^]_i_ to near-diastolic levels).^8-10^ As a result of RRC dyscoupling, a potentially vulnerable period for MAR (VP_MAR_) arises in late systole, which scales with T_RRC_, as high [Ca^2+^]_i_ favours arrhythmogenesis in progressively re-excitable CM.

In the present study, rabbit isolated left ventricular (LV) CM were subjected to acute stretch using a carbon fibre (CF) system,^6^ combined with video-based measurement of sarcomere length dynamics, simultaneous fluorescence imaging of V_m_ and [Ca^2+^]_i_, and pharmacological interrogations, to determine mechanisms underlying systolic MAR in CM with an ischaemia- or pharmacologically-induced increase in the duration of VP_MAR_.

## METHODS

### Data Availability

Key methodological information is provided below, with details found in the Supplemental Material. The data and computer code that support the findings of this study are available from the corresponding author upon reasonable request.

### Ethical Approval

Experiments were conducted in accordance with the ethical guidelines of the Canadian Council on Animal Care and the German legislation for animal welfare, following protocols approved by the Dalhousie University Committee for Laboratory Animals and the local Institutional Animal Care and Use Committee at the University of Freiburg. Experimental methodologies followed the Guidelines for Assessment of Cardiac Electrophysiology and Arrhythmias in Small Animals,^11^ with details reported following the Minimum Information about a Cardiac Electrophysiology Experiment (MICEE) reporting standard.^12^

### Langendorff-based Isolated CM Preparation

For experiments involving cell stretch or fluorescence imaging, LV CM were isolated from hearts of female New Zealand White rabbits (2.1 ± 0.2 kg, Charles River), and exposed to: (i) control solution (CTRL; containing, in mM: 142 NaCl, 4.7 KCl, 1 MgCl_2_, 1.8 CaCl_2_, 10 glucose, and 10 HEPES [Sigma-Aldrich]; pH adjusted to 7.40 ± 0.05 with NaOH; osmolality 300 ± 5 mOsm/L); (ii) a simulated ischaemia (SI) solution with a composition that mimics extracellular fluid after 30 min of ischaemia^13,14^ (containing, in mM: 140 NaCl, 15 KCl, 1.8 CaCl_2_, 1 MgCl_2_, 10 HEPES, 1 NaCN to block oxidative phosphorylation, and 20 2-deoxyglucose to block anaerobic glycolysis [Sigma-Aldrich]; pH adjusted to 6.5 ± 0.05 with NaOH, osmolality 300 ± 5 mOsm/L); or (iii) CTRL solution containing pinacidil (PIN; 50 µM, to activate ATP-sensitive potassium [K_ATP_] channels).

### CF-based Single CM Stretch

Single CM were stretched using the CF method, adapted from previous work.^6,13^ Acute stretch, approximating paradoxical segment lengthening of ischaemically weakened myocardium during regional ischaemia,^15^ was applied to the cell (20 µm piezo-electric actuator displacement, applied and removed at a rate of 0.7 μm/ms, with a total stretch and release pulse duration of 110 ms; Figure 1a, b). Stretch was timed from the electrical stimulus to commence during mid-diastole (600 ms delay) or during the VP_MAR_ (300 ms delay in CTRL, 150 ms during exposure to SI or PIN, and 210 ms during exposure to SI or PIN with glibenclamide, based on the timing measured by simultaneous V_m_-Ca^2+^ imaging, described below), which was repeated with 10 s between each stretch, for a total of four stretches (Figure 1b). This protocol was repeated with increasing magnitudes of displacement (30 µm and 40 µm, with total stretch and release pulse durations of 140 ms or 170 ms) to generate a range of stretch-induced changes in sarcomere length within a CM (30 s between repetitions, for a total of 12 stretches; Figure 1b). Sarcomere length and the piezo-electric actuator and CF tip positions were monitored and recorded at 240 Hz (Myocyte Contractility Recording System, IonOptix), which were used to measure CM contractile function, characteristics of CM stretch, and the incidence of MAR.

**Figure 1.**
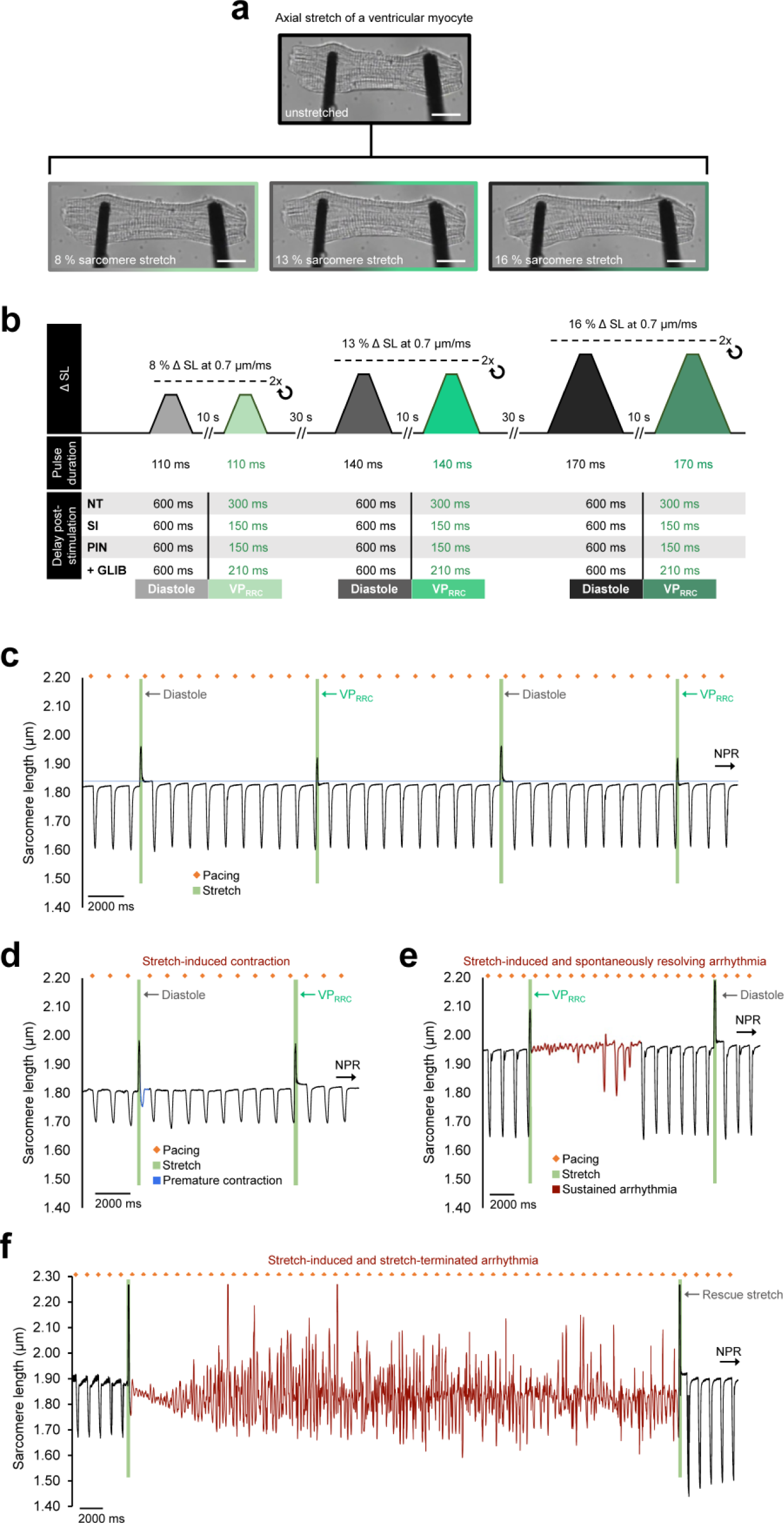
Arrhythmias elicited by acute stretch of rabbit left ventricular cardiomyocytes (LV CM). **a,** Brightfield image of a rabbit LV CM before (top) and during (bottom) axial stretch, applied using a carbon fibre-based system. Scale bar: 10 μm. **b**, Schematic of the stretch protocol. **c**, Representative measurement of sarcomere length in a CM during 1 Hz pacing (orange dots) and stretch (green segments) applied in diastole (stretches 1 and 3) or in late systole (stretches 2 and 4), which did not result in an arrhythmia (normal paced rhythm, NPR). **d,** Stretch-induced contraction (blue curve segment) upon diastole-timed stretch. **e,** Arrhythmic activity (red segment) after a stretch timed to the vulnerable period for mechano-arrhythmogenesis (VP_MAR_) that spontaneously resolved. **f,** Sustained arrhythmic activity (red segment) after a diastolic stretch that was terminated by application of an additional stretch.

### Pharmacological Interventions

Mechanisms contributing to MAR were assessed by pre-treating CM with pharmacological agents, including: glibenclamide (to block K_ATP_ channels; 20 μM for 15 min, from a 10 mM stock in DMSO, Abcam); BAPTA-AM (to buffer [Ca^2+^]_i_; 1 µM for 20 min, from a 10 mM stock in dimethyl sulfoxide [DMSO], Abcam, with the concentration determined in preliminary experiments by titrating to a value that caused a ∼10 % decrease in percent sarcomere shortening during contraction); dantrolene (to stabilise ryanodine receptors; 1 µM for 5 min, from a 5 mM stock in DMSO, Abcam); streptomycin (to non-specifically block mechano-sensitive channels; 50 µM for 5 min, from a stock concentration of 15 mM in distilled water, Sigma-Aldrich); allyl isothiocyanate (AITC, to activate transient receptor potential kinase ankyrin 1 [TRPA1] channels; 10 µM for 5 min, from a stock concentration of 1 M in DMSO, Sigma-Aldrich); HC-030031 (to block TRPA1 channels; 10 µM for 30 min, from a 50 mM stock in DMSO, Abcam), N-acetyl-L-cysteine (NAC, to chelate reactive oxygen species [ROS]; 10 mM for 20 min, from a 0.5 M stock in distilled water, Sigma-Aldrich); and diphenyleneiodonium (DPI, to block ROS production; 3 µM for 60 min, from a 10 mM stock in DMSO, Abcam).

### Simultaneous V_m_-Ca^2+^ Fluorescence Imaging

The CM were dual-stained with V_m_ and [Ca^2+^]_i_-sensitive fluorescent dyes (di-4-ANBDQPQ and Fluo-5F-AM) and simultaneous measurement of V_m_ and Ca^2+^ was performed by single-excitation, dual-emission fluorescence imaging with a single camera, optical splitter system (Figure 2a).^16^

**Figure 2.**
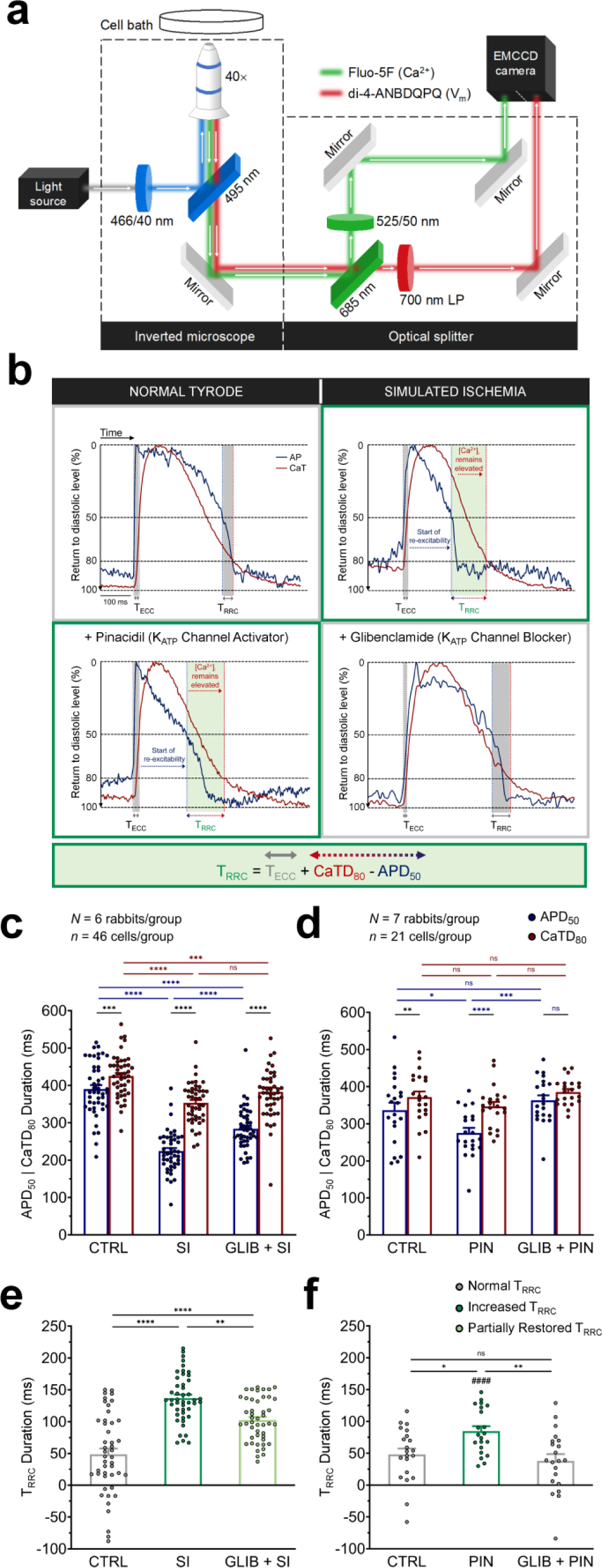
Dyscoupling of repolarisation-relaxation coupling (RRC) in rabbit left ventricular cardiomyocytes (LV CM) by simulated ischaemia (SI) or ATP-sensitive potassium (K_ATP_) channel activation with pinacidil (PIN). **a**, Schematic of the single-excitation, dual-emission fluorescence imaging approach, utilising a single camera with an image splitter for simultaneous measurement of voltage and calcium (Ca^2+^) in rabbit LV CM. **b,** Representative signals showing the duration of RRC (T_RRC_, grey) in control (CTRL, top left), after dyscoupling of RRC with SI (top right) or PIN (bottom left), both leading to increased T_RRC_ (green), and upon mitigation of RRC dyscoupling in SI by application of glibenclamide (GLIB, bottom right), decreasing in T_RRC_ closer to CTRL. **c,** Measurements of action potential duration at 50 % repolarisation (APD_50_, blue) and Ca^2+^ transient duration at 80 % return to diastolic levels (CaTD_80_, red) in CTRL, SI, or SI following pre-treatment with GLIB (GLIB + SI). **d,** Measurements during CTRL or exposure to PIN, or GLIB + PIN. **e,f,** T_RRC_ (= excitation-contraction coupling duration [T_ECC_] + CaTD_80_ - APD_50_), in each condition. Error bars represent standard error of the mean. Differences between conditions assessed by one-way ANOVA, with Tukey *post-hoc* tests, and between APD_50_ and CaTD_80_ or SI and PIN by paired, two-tailed, Student’s *t*-tests. \**p* < 0.05, \*\**p* < 0.01, \*\*\**p* < 0.001, and \*\*\*\**p* < 0.0001. ^####^*p* < 0.0001 for SI *vs* PIN. *N* = rabbits / group, *n* = cells / group.

### Ratiometric [Ca^2+^]_i_ Fluorescence Imaging

The CM were stained with a ratiometric [Ca^2+^]_i_-sensitive fluorescent dye (Fura Red AM) and measurement of [Ca^2+^]_i_ levels was performed by near-simultaneous collection of ratiometric signals using a dual-excitation, single-emission imaging with frame-synchronised, pulsed-excitation approach.^17^

### Tissue Slice-based Isolated CM Preparation

For experiments involving patch clamp recordings of TRPA1 current, LV CM were isolated from slices of LV myocardium from hearts of female New Zealand White rabbits (2.0 ± 0.2 kg, Charles River) according to previously published protocols.^18,19^

### Sarcolemmal Ion Current Recordings

Sarcolemmal ion current was recorded by the patch-clamp technique in cell-attached configuration with protocols previously used to study cation non-selective channels.^20^

### Statistical Analysis

Values are reported as mean±SEM. Statistical tests were performed using Prism 9 (GraphPad), with significance indicated by *p* < 0.05. Applied tests are indicated in the figure captions, along with the number of replicates.

## RESULTS

### Acute Stretch Triggers MAR in CM

Acute stretch (with a total stretch-and-release duration of 110-170 ms, to mimic paradoxical segment lengthening as occurs in ischaemically weakened myocardium in the intact heart)^15^ was applied with CF, attached to both ends of single LV CM under piezo-electric actuator control (Figure 1a). Stretch (timed with a variable delay from the electrical pacing stimulus to occur during diastole or late systole, Figure 1b) resulted in a variety of disturbances in the rhythm of CM mechanical activity (revealed by tracking sarcomere length, Figure 1c), including stretch-induced contractions (sometimes followed by a second contraction out of sync with the continuing electrical stimulation; Figure 1d and Video S1) and sustained rhythm disturbances (3 or more successive contractions immediately after stretch), which either resolved spontaneously (Figure 1e) or were terminated by application of an additional stretch (Figure 1f, Video S2).

Since previous work in the intact heart has shown that the magnitude of tissue deformation is a key determinant of MAR,^21^ whether the incidence of MAR in CM is also dependent on stretch amplitude was assessed. Increasing piezo-electric actuator displacement from 20 to 40 μm resulted in an increase in the maximum sarcomere length during stretch, the percent sarcomere stretch, and the maximum applied stress in all CM (Figures 1a and S1). The increase in applied stretch corresponded with an increased incidence of rhythm disturbances (Figure S2), illustrating that MAR in CM, as in the intact heart, is dependent on stretch magnitude.

### K_ATP_ Channel Activation Dyscouples RRC

To replicate ischaemic changes in AP morphology, LV CM were exposed to: (i) SI solution, which combined metabolic inhibition (block of oxidative phosphorylation and anaerobic glycolysis) with acidosis and hyperkalemia to mimic environmental conditions at 30 min of ischaemia,^13,14^ or (ii) PIN-containing solution, which is a well-established agonist of sulfonylurea receptor 2A / K_ir_6.2 (the hetero-octamer that forms the K_ATP_ channel in cardiac and skeletal muscle^22^), to assess the isolated effects of ischaemia-induced K_ATP_ channel opening. To measure the effects of these interventions on RRC, single-excitation, dual-emission fluorescence imaging was performed. The dyes di-4-ANBDQPQ and Fluo-5F AM (whose relatively high K_d_ value [∼2.3 μM] limits its buffering of diastolic [Ca^2+^]_i_ and thus helps to avoid potential artefactual lengthening of Ca^2+^ transient duration, CaTD) were used to simultaneously measure V_m_ and [Ca^2+^]_i_ transients in electrically-paced rabbit LV CM under CF control (Figure 2a). T_RRC_ was quantified as the time difference in the post-activation recovery of [Ca^2+^]_i_ and V_m_ (calculated as ECC duration [T_ECC_] plus CaTD at 80 % return to diastolic levels [CaTD_80_], minus AP duration at 50 % repolarisation [APD_50_]), which is a period during which CM begin to become re-excitable while [Ca^2+^]_i_ remains elevated (Figure 2b).

Under physiological conditions with normal RRC (using CTRL solution), CaTD_80_ was significantly greater than APD_50_ (individually true in the majority of cases; Figure 2b-d). During exposure to SI or PIN, APD_50_ (Figure 2b-d) decreased. CaTD_80_ also decreased during SI (which can be expected with metabolic inhibition and acidosis^24^), but to a lesser extent than APD_50_, whereas with PIN there was no significant change in CaTD_80_. As a result, there was dyscoupling of RRC both with SI and with PIN, which resulted in an increase in T_RRC_ (Figure 2b,e-f; with a greater increase in SI compared to PIN). These changes in APD_50_, CaTD_80_, and T_RRC_ mimic behaviour previously reported in intact hearts during acute ischaemia^8,9^ and in hypoxic conditions.^10^ In CM pre-treated with glibenclamide (to block K_ATP_ channels), the reduction in APD_50_ was attenuated (during SI) or prevented (with PIN), with no significant change in CaTD_80_ (Figure 2b-d), such that T_RRC_ decreased compared to either SI or PIN in the absence of glibenclamide (Figure 2b,e,f).

### RRC Dyscoupling Permits Systolic MAR

To determine whether an increase in T_RRC_ affects the incidence of systolic MAR, stretch protocols were repeated with CM exposed to SI or to PIN. In both cases, there was an increase in the incidence of MAR with stretch in late systole compared to CTRL (Figure 3a) and, as in CTRL, an increase in stretch magnitude (quantified as increased percent sarcomere stretch, maximum sarcomere length during stretch, and maximum applied stress; Figure S1) raised the incidence of rhythm disturbances (Figure S2). Importantly, there was no difference in the percent sarcomere stretch, maximum sarcomere length during stretch, or maximum applied stress across groups (CTRL, SI, or PIN; Figure S1), indicating that there were no changes in passive mechanical properties of the CM, or subsequent changes in the characteristics of the stretch or stress they experienced, that could account for the increase in MAR during SI or PIN.

**Figure 3.**
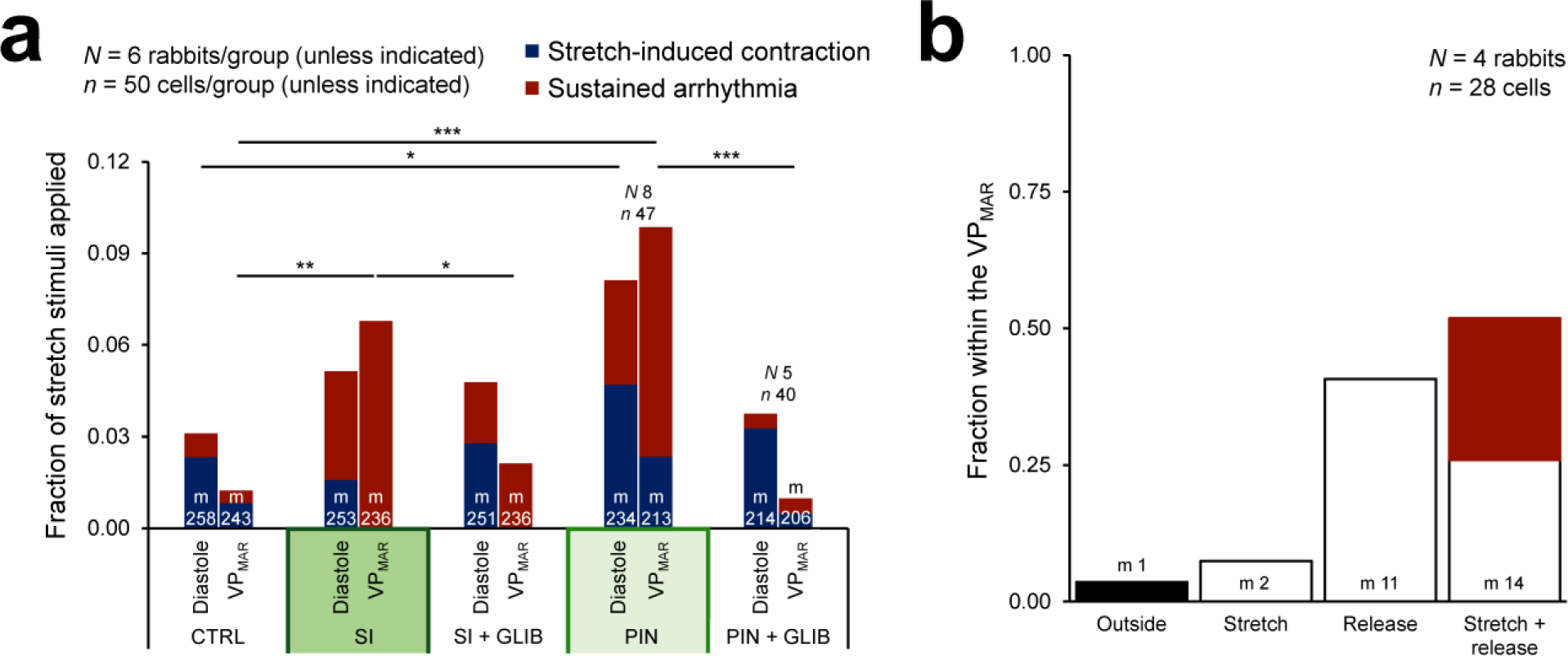
Role of repolarisation-relaxation coupling (RRC) dyscoupling in mechano-arrhythmogenesis (MAR). **a,** Incidence of stretch-induced contractions (blue) and sustained arrhythmias (red) with stretch of rabbit left ventricular cardiomyocytes (LV CM) during diastole or the vulnerable period for MAR (VP_MAR_) in control (CTRL), or during exposure to simulated ischaemia (SI), glibenclamide (GLIB) + SI, pinacidil (PIN), or GLIB + PIN. **b,** Fraction of phases of stretch stimuli (stretch, release, stretch + release) within the VP_MAR_, with those that resulted in arrhythmias shown in red. Differences in arrhythmia incidence assessed using chi-square contingency tables and Fisher’s exact test. \**p* < 0.05, \*\**p* < 0.01, and \*\*\**p* < 0.001 between groups. *N* = rabbits / group, *n* = cells / group, m = stretch stimuli applied.

To test whether SI- or PIN-induced dyscoupling of RRC, and thus an increase in the duration of VP_MAR_, contributed to the increased probability of systolic MAR, CM were pre-treated with glibenclamide. Glibenclamide prevented the increase in systolic MAR in both cases (Figure 3a), suggesting that the presence of physiological RRC protects CM from arrhythmogenesis, while RCC dyscoupling and an increase in VP_MAR_ duration raise MAR susceptibility during late phases of the AP. Of note, glibenclamide pre-treatment had no significant effect on diastolic MAR in CM exposed to either SI or PIN (Figure 3a).

SI had no significant effect on the incidence of MAR with stretch in diastole compared to CTRL, indicating that ischaemic dyscoupling of RRC does not affect diastolic MAR. In contrast, there was an increased incidence of rhythm disturbances with diastolic stretch in PIN-treated CM (Figure 3a). This surprising result may be related to secondary effects of PIN, which were explored in additional experiments, described in the following section.

To ascertain whether stretch-induced cell damage could account for the observed increases in MAR, CM contractile function was assessed before and after each magnitude of stretch. In CM exposed to CTRL, SI, or PIN, diastolic sarcomere length and the maximum rate and percent of sarcomere shortening were not significantly different before and after stretch application, suggesting that CM damage or CF slippage during stretch are unlikely to have occurred (Figure S3). Recorded parameters were also not significantly different before and after a return to steady state following periods of sustained rhythm disturbances (Figure S4), indicating that cellular damage is also unlikely to underlie the more severe forms of MAR.

### [Ca^2+^]_i_ Plays a Role in MAR in CM

During ischaemia, altered Ca^2+^ handling results in elevated [Ca^2+^]_i_,^23^ which can be arrhythmogenic by driving forward-mode sodium / Ca^2+^-exchanger (NCX) activity, and by shifting CM V_m_ closer to the threshold for excitation.^24^ This may facilitate systolic MAR, as an influx of Ca^2+^ through mechano-sensitive ion channels during stretch will further drive NCX-mediated depolarisation, while at the same time directly cause V_m_ depolarisation,^5^ which can trigger arrhythmic electrical activity.

The importance of [Ca^2+^]_i_ for MAR during ischaemia has been demonstrated in the intact heart.^8^ To assess whether [Ca^2+^]_i_ also plays a role in CM-level MAR, [Ca^2+^]_i_ was buffered with BAPTA-AM (1 μM, K_d_ ≍ 0.5 μM), which has previously been shown in ventricular CM to prevent stretch-induced changes in cardiac electrophysiology driven by increases in [Ca^2+^]_i_.^25^ In CM with SI- or PIN-induced dyscoupling of RRC, BAPTA-AM reduced the incidence of systolic MAR (Figure 4a; in CTRL, BAPTA-AM application had no significant effect, Figure S5). Ca^2+^ buffering also prevented the increase in diastolic MAR that occurs in PIN-treated cells (Figure 4a). Surprisingly, though, BAPTA-AM increased the incidence of diastolic MAR in CM exposed to SI (Figure 4a), in particular of single stretch-induced beats, which may be explained by a Ca^2+^ buffering-induced increase in the trans-sarcolemmal driving force for Ca^2+^ entry during stretch, specifically in ischaemic conditions.

**Figure 4.**
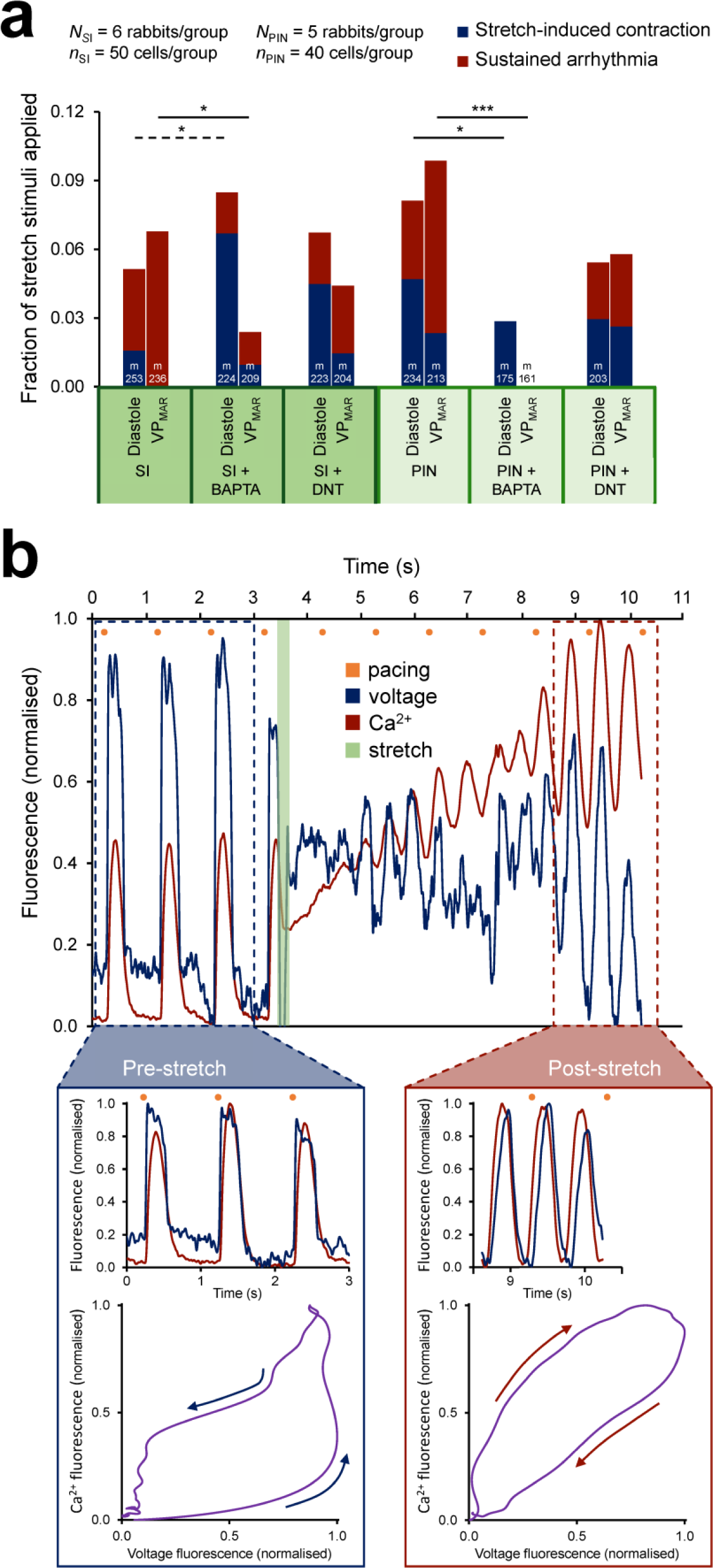
Contribution of cytosolic calcium concentration ([Ca^2+^]_i_) to systolic mechano-arrhythmogenesis (MAR). **a,** Incidence of stretch-induced contractions (blue) and sustained arrhythmias (red) with stretch of rabbit left ventricular cardiomyocytes (LV CM) during diastole or the vulnerable period for MAR (VP_MAR_) during exposure to simulated ischaemia (SI) or pinacidil (PIN), as well as in combination with BAPTA-AM (to buffer [Ca^2+^]_i_) or dantrolene (DNT, to stabilise ryanodine receptors in their closed state). **b,** Representative voltage (blue) and Ca^2+^ (red) signals simultaneously recorded by fluorescence imaging in a CM exposed to PIN, showing a sustained arrhythmia induced by stretch during the VP_MAR_ (green), with persistent depolarisation and an associated increase in [Ca^2+^]_i_ above normal systolic levels, followed by spontaneous oscillations in [Ca^2+^]_i_ (which subsequently resolved, followed by normal paced beats; not shown). **Blue inset**: Scaled voltage and Ca^2+^ signals (top) and phase plot of the first beat (bottom), showing that prior to stretch, changes in voltage preceded Ca^2+^. **Red inset**: Scaled voltage and Ca^2+^ signals (top) and phase plot of the first beat (bottom), showing [Ca^2+^]_i_ oscillations preceding changes in voltage during the stretch-induced sustained arrhythmia, suggesting aberrant Ca^2+^ handling as the driving mechanism. Differences in arrhythmia incidence assessed using chi-square contingency tables and Fisher’s exact test. \**p* < 0.05 and \*\*\**p* < 0.001 between groups (dashed line indicates an increase). *N* = rabbits / group, *n* = cells / group.

As stretch is also known to increase Ca^2+^ leak from the sarcoplasmic reticulum through ryanodine receptors,^6^ which is enhanced in ischaemia^13^ and may contribute to the associated increase in [Ca^2+^]_i_,^26^ a stretch-induced increase in Ca^2+^ leak may also contribute to the ischaemia-induced increase in systolic MAR. To investigate this possibility, dantrolene was used to stabilise ryanodine receptors in their closed state, which has been shown to not affect Ca^2+^-induced Ca^2+^ release or contractile function of CM.^27^ Dantrolene had no significant effect on the incidence of MAR in CM exposed to SI or PIN (Figure 4a; in CTRL there was also no significant effect, Figure S5).

To assess whether the reduction in MAR with BAPTA-AM was a result of secondary effects on CM mechanics, contractile function was assessed in CTRL cells. Treatment with BAPTA-AM (as well as dantrolene) had no effect on diastolic sarcomere length or maximum rate of systolic sarcomere shortening, although there was a slight decrease in the percent systolic sarcomere shortening with BAPTA-AM, likely due to buffering of peak systolic [Ca^2+^]_i_ (Figure S6).

The above findings suggest that [Ca^2+^]_i_ is important for MAR in CM with RRC dyscoupling. The most definitive evidence for a direct role of [Ca^2+^]_i_ in MAR, though, came from dual V_m_-Ca^2+^ fluorescence imaging of CM during sustained rhythm disturbances. This revealed progressive increases in [Ca^2+^]_i_ after stretch application, which were followed by Ca^2+^ oscillations that preceded changes in V_m_, indicating that [Ca^2+^]_i_ fluctuations can sustain stretch-triggered arrhythmias (Figure 4b, Video S3).

### Systolic MAR Depends on Stretch and Release

The results to this point demonstrate an increase in Ca^2+^-mediated systolic MAR in CM with dyscoupling of RRC. While arrhythmias appear to occur with stretch during the VP_MAR_, the termination of stretch (‘release’) may also be arrhythmogenic. Stretch of cardiac tissue increases the affinity of myofilaments for Ca^2+^, such that stretch during systole, when [Ca^2+^]_i_ is high, results in increased Ca^2+^ binding. Upon release, rapid dissociation of myofilament-bound Ca^2+^ may produce a surge in [Ca^2+^]_i_, which triggers Ca^2+^ release and generates Ca^2+^ waves or drives NCX-mediated membrane depolarisation.^7^

In a subset of CM exposed to SI, fluorescence-based measurements of APD and CaTD during late-systolic stretch was performed to assess the phases of the stretch pulse (*i.e.,* stretch, release, or stretch-and-release) that occurred during the VP_MAR_, and relate this to the effects of the mechanical stimulus on systolic MAR. The analysis confirmed that in 96 % of CM, at least part of the stretch pulse was timed to occur during the CM-specific VP_MAR_. MAR was induced only if both stretch *and* release occurred during the VP_MAR_, which was true in just over half of the cells tested. In these CM, MAR was triggered in 50 % of cases (Figure 3b). These results suggest that the combination of stretch-and-release may be particularly prone to causing late-systolic MAR. They also confirm targeting of mechanical stimulation to the VP_MAR_ in the experiments presented above, and may account for the lower incidence of MAR seen in those experiments (Figure 3 and 4), as it appears that in only ∼50% of cases stretch-and-release occurs during the VP_MAR_ and only ∼50% of those will result in MAR.

### Activated TRPA1 Channels Drive MAR

While it was anticipated that RRC dyscoupling in CM during exposure to SI or to PIN would raise the incidence of systolic MAR, the increase in diastolic MAR with PIN (Figure 3a), and what appears to be a trend toward a more pronounced increase in systolic MAR with PIN compared to SI (Figure 3a; on the background of a less pronounced increase in T_RRC_, Figure 2e-f), was surprising. Thus, experiments were performed to determine mechanisms underlying the overall increase in MAR with PIN.

PIN is generally considered a specific agonist of K_ATP_ channels, resulting in a repolarising efflux of potassium from CM. However, in HEK293 cells, PIN has been shown to increase the activity of TRPA1 channels, resulting in a depolarising influx of Ca^2+^.^28^ This is potentially important in the context of MAR, as TRPA1 channels: (i) are inherently mechano-sensitive^29^ and contribute to mechanically-evoked electrical responses (measured as trans-membrane currents and deduced from AP initiation) in sensory neurons,^30-33^ astrocytes,^34^ and vertebrate hair cells;^35^ (ii) are functionally expressed in murine ventricular CM^36^ and contribute to changes in *Drosophila* heart rhythm in response to mechanical stimulation;^37^ and (iii) are important for cardiovascular function and disease progression.^38^ It was hypothesised that if TRPA1 channels are functionally expressed in rabbit LV CM, they could contribute to the increase in systolic and diastolic MAR upon PIN treatment.

To evaluate the functional expression of TRPA1 channels in rabbit LV CM, ion channel activity was measured in cell-attached patches during application of the TRPA1 channel-specific agonist, AITC.^39^ AITC exposure caused an increase in total inward current, (with a ∼1-3 min delay, similar to the time-lag in response reported by others^40-41^), while there was no change in time-matched CTRL cells (Figure 5a, b). To explore whether activated TRPA1 channels can contribute to MAR, CM were exposed to AITC (which has been shown to also enhance the response of TRPA1 channels to mechanical stimulation^32^) and subjected to diastolic stretch. CM exposed to AITC showed increased diastolic MAR compared to CTRL, which was prevented by application of BAPTA-AM, as well as by application of the specific TRPA1 channel antagonist HC-030031^42^ (which has been shown to inhibit TRPA1 current and mechanically-induced excitation in sensory neurons;^30-33^ Figure 5c).

**Figure 5.**
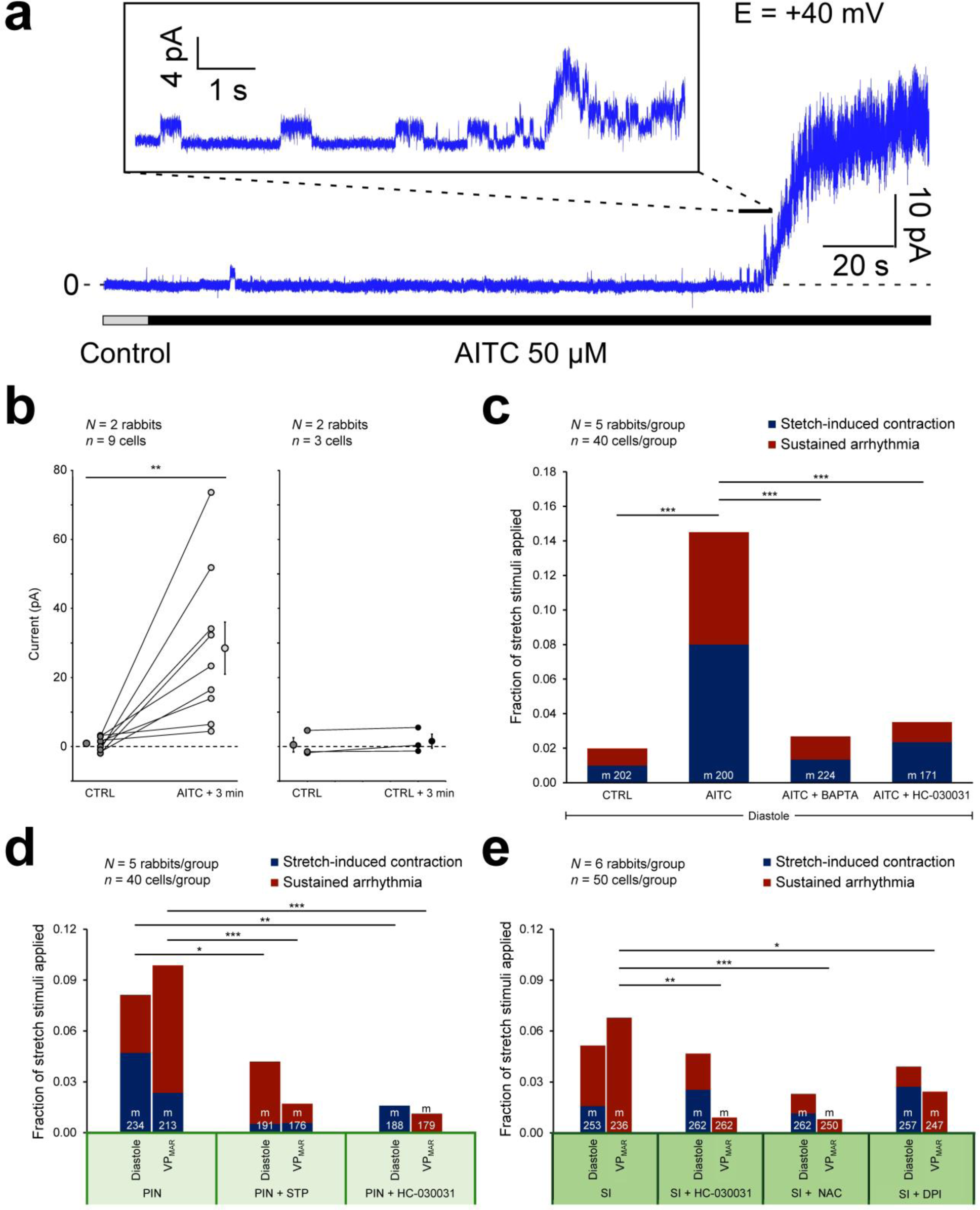
Contribution of transient receptor potential kinase ankyrin 1 (TRPA1) channels to mechano-arrhythmogenesis (MAR) during dyscoupling of repolarisation-relaxation coupling (RRC). **a,** Representative current recording by cell-attached patch (holding potential +40 mV) in rabbit left ventricular cardiomyocytes (LV CM) in control (CTRL) and after switching to allyl isothiocyanate (AITC)-containing solution (50 μM; to activate TRPA1 channels). **Inset**, Detail of channel activation with visible single channel events. **b**, Quantification of AITC-induced current changes (left; 3 min total application with a 15 s lag time for AITC to reach the CM) and time-matched, same-batch CTRL CM (right; 3 min). Error bars represent standard error of the mean. **c,** Incidence of stretch-induced contractions (blue) and sustained arrhythmias (red) with stretch of CM during diastole in CTRL, or during exposure to AITC, AITC + BAPTA-AM (to buffer cytosolic calcium concentration, [Ca^2+^]_i_), or AITC + HC-030031 (to block TRPA1 channels). **d**, Incidence of MAR with stretch of CM during diastole or the vulnerable period for MAR (VP_MAR_) during exposure to pinacidil (PIN), PIN + streptomycin (STP, a non-specific blocker of mechano-sensitive ion channels), or PIN + HC-030031. **e**, Incidence of MAR during exposure to simulated ischaemia (SI), or to SI + HC-030031, N-acetyl-L-cysteine (NAC, to chelate reactive oxygen species, ROS), or diphenyleneiodonium (DPI, to block ROS production). Current differences assessed with two-tailed, paired, Student’s *t*-test. \*\**p* < 0.01. Differences in arrhythmia incidence assessed using chi-square contingency tables and Fisher’s exact test. \**p* < 0.05, \*\**p* < 0.01, and \*\*\**p* < 0.001 between groups. *N* = rabbits / group, *n* = cells / group, m = stretch stimuli applied.

Having demonstrated that TRPA1 channels are functionally expressed and appear to be a source of Ca^2+^-mediated MAR in rabbit LV CM, whether TRPA1 channels contribute to the increase in systolic and diastolic MAR with PIN was assessed. LV CM were subjected to PIN, or PIN with either streptomycin (a non-specific antagonist of cation non-selective mechano-sensitive ion channels^43^) or with the TRPA1-specific antagonist HC-030031 and subjected to stretch in diastole or during the VP_MAR_. Streptomycin and HC-030031 both reduced the incidence of MAR with diastolic and late systolic stretch in PIN-treated CM (Figure 5d; MAR incidence remained low in CTRL [Figure S5], and neither drug had any other significant effect on functional assays [Figure S6]), supporting the suggestion that TRPA1 channels may contribute to the increased incidence of both diastolic and systolic MAR caused by PIN.

As TRPA1 channels are known to be activated by ischaemic conditions,^44^ whether TRPA1 channels are also involved in the increase in systolic MAR seen during SI was investigated. It was found that blocking TRPA1 channels with HC-030031 prevented the increase in systolic MAR during SI, suggesting their involvement in that setting as well (Figure 5e).

To assess whether the reduction in MAR with streptomycin or HC-030031 and the increase with AITC might have been the result of secondary effects on CM mechanics, contractile function after drug treatment was assessed. In all cases there was no significant effect on diastolic sarcomere length or maximum rate or percent of systolic sarcomere shortening (Figure S6).

### ROS Mediates MAR in Ischaemia

TRPA1 channels are known to be activated by an increase in [Ca^2+^]_i_^45^ or ROS,^46^ both of which are increased during ischaemia.^23^ Experiments using BAPTA-AM to buffer [Ca^2+^]_i_ suggested a contribution of [Ca^2+^]_i_ to systolic MAR in SI, which may in part involve activation of TRPA1. To determine whether ROS also contributes to the ischaemia-induced increase in systolic MAR, prior to SI, CM were incubated with NAC (to chelate intracellular ROS) or DPI (to inhibit NOX2-mediated mechano-sensitive ROS production, which is enhanced in ischaemia^13^). The incidence of systolic MAR was decreased by both NAC and DPI (Figure 5e), suggesting a contribution of ROS to systolic MAR during SI. The contribution of ROS to MAR may act *via* a ROS-mediated increase in TRPA1 activity (enhancing its mechano-sensitive depolarising current, as well as its contribution to increased [Ca^2+^]_i_^47^), or through a TRPA1-independent increase in [Ca^2+^]_i_.^48^

### TRPA1 and [Ca^2+^]_i_ Can Each Drive MAR

Combined, the above results suggest that increased mechano-sensitive TRPA1 channel activity and increased [Ca^2+^]_i_ contribute to a higher incidence of MAR during SI and PIN-induced K_ATP_ activation. However, as it had been shown that TRPA1 itself can lead to increased [Ca^2+^]_i_,^47^ it remained unclear whether the two mechanisms may independently contribute to MAR. [Ca^2+^]_i_ was measured by ratiometric fluorescence imaging, which showed that diastolic [Ca^2+^]_i_ increased during exposure to SI, PIN, or AITC. This increase, however, was reduced by TRPA1 channel block with HC-030031 only during exposure to AITC (Figure S7), suggesting that TRPA1 channels are the primary driver of an increase in [Ca^2+^]_i_ during activation with AITC, while with SI or PIN the increase is driven by other mechanisms (Figure S7; of note: in SI, chelating ROS with NAC also did not prevent the increase in diastolic [Ca^2+^]_i_). As blocking of TRPA1 channels with HC-030031 (Figure 5c-e) and buffering of [Ca^2+^]_i_ with BAPTA-AM (Figure 4a, 5c) decreased MAR during exposure to SI, PIN, and AITC, it would appear that increased TRPA1 activity and increased [Ca^2+^]_i_ content can independently contribute to increased MAR (unfortunately, the effects of BAPTA-AM on [Ca^2+^]_i_ could not be measured, as the combined buffering of Ca^2+^ by BAPTA-AM and Fura Red AM decreased CM contractile function).

## DISCUSSION

Taken together, the results from this study demonstrate that dyscoupling of RRC in LV CM during ischaemia or by pharmacological activation of K_ATP_ channels (to mimic their opening in ischaemia) results in an increase in the incidence of systolic MAR, while normal RRC or prevention of RRC dyscoupling by K_ATP_ channel antagonism are protective. Key molecular mechanisms driving an ischaemia-induced increase in systolic MAR are activation of TRPA1 channels and an increase in [Ca^2+^]_i_, both of which can be exacerbated by an increase in ROS. Figure 6 summarises key findings and presents a conceptual model of the contribution of RRC dyscoupling, TRPA1 channels, [Ca^2+^]_i_, and ROS to the triggering (*via* a depolarising trans-sarcolemmal current) and sustenance (*via* the generation of a [Ca^2+^]_i_-mediated substrate) of systolic MAR in ischaemic LV CM.

**Figure 6.**
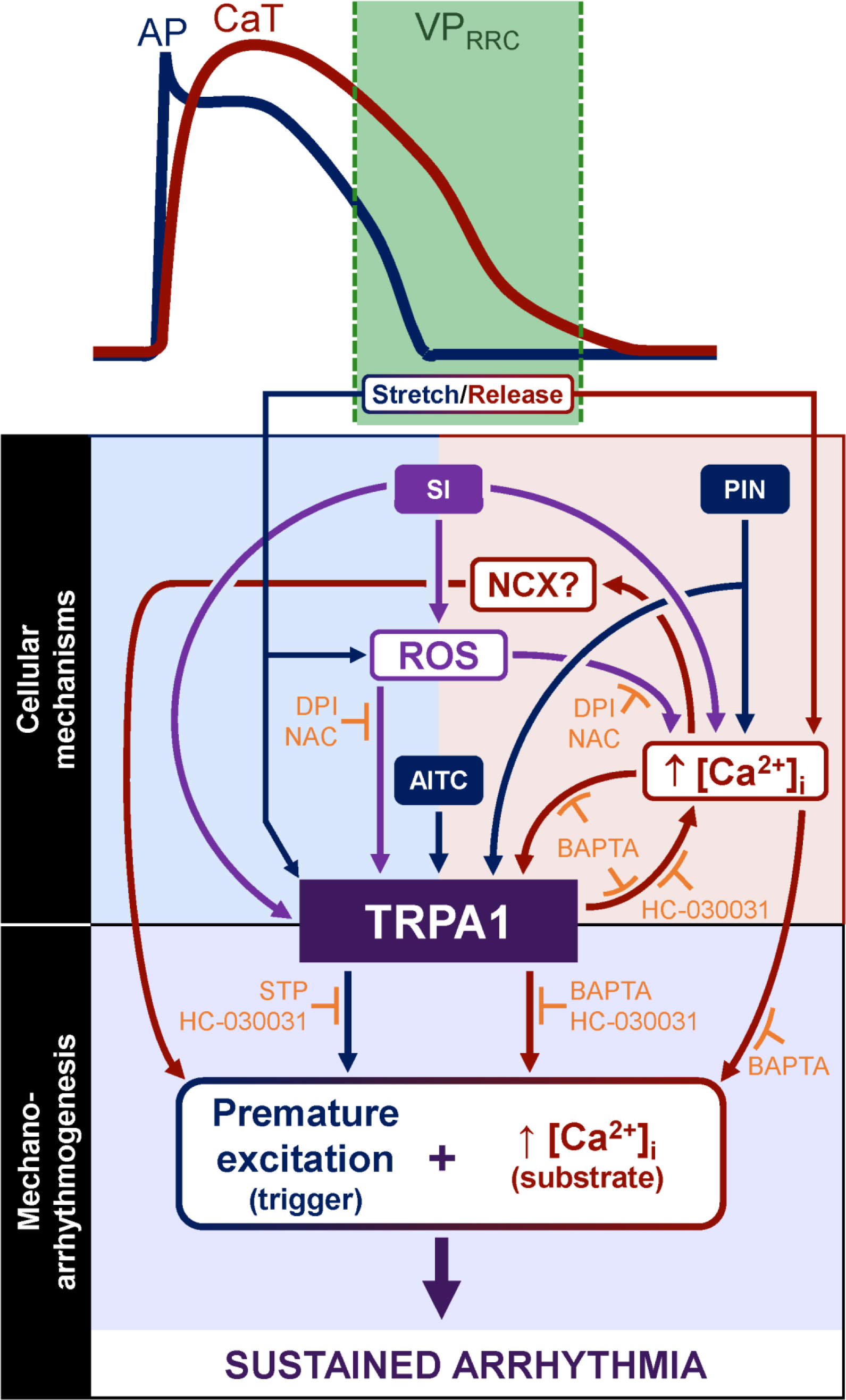
Role of repolarisation-relaxation coupling (RRC) dyscoupling, transient receptor potential kinase ankyrin 1 (TRPA1) channels, cytosolic calcium concentration ([Ca^2+^]_i_), and reactive oxygen species (ROS) in ventricular systolic mechano-arrhythmogenesis (MAR). Schematic of the proposed mechanisms underlying the TRPA1- and Ca^2+^-mediated increase in systolic MAR during dyscoupling of RRC. AITC, Allyl isothiocyanate; AP, action potential; CaT, Ca^2+^ transient; DNT, dantrolene; DPI, diphenyleneiodonium; NAC, N-acetyl-L-cysteine; NCX, sodium-Ca^2+^ exchanger; PIN, pinacidil; ROS, reactive oxygen species; SI, simulated ischaemia; STP, streptomycin; VP_MAR_, vulnerable period for mechano-arrhythmogenesis.

As indicated in the figure, triggered activity during ischaemia may be driven by mechano-sensitive TRPA1 channels.^29^ Stretch is known to increase the trans-sarcolemmal influx of cations through activated TRPA1 channels (which preferentially pass Ca^2+^ in HEK293 cells^49^), directly leading to V_m_ depolarisation.^30-35^ At the same time, the resulting increase in [Ca^2+^]_i_, along with a dissociation of myofilament-bound Ca^2+^ during the release of stretch,^7^ may cause V_m_ depolarisation *via* electrogenic forward-mode NCX activity.^48^ If the resulting V_m_ depolarisation is sufficiently large, it will trigger CM excitation and contraction. This is particularly arrhythmogenic when ischaemic dyscoupling of RRC allows both stretch and release to occur during the VP_MAR_ (the period during which [Ca^2+^]_i_ remains high in progressively re-excitable CM). The resulting triggered activity may devolve into sustained arrhythmias if [Ca^2+^]_i_ stays sufficiently elevated, creating an arrhythmogenic substrate.^24^ In addition, an increase in [Ca^2+^]_i_ will further increase TRPA1 channel activity, as it is directly modulated by [Ca^2+^]_i_^46^ (although the level of TRPA1 activation will be limited as [Ca^2+^]_i_ increases, by simultaneous Ca^2+^-mediated inactivation^50^).

It was found that exposure to AITC caused direct activation of TRPA1 channels and subsequent diastolic [Ca^2+^]_i_ loading, while PIN either directly activated TRPA1 channels or did so indirectly *via* an increase in [Ca^2+^]_i_. During SI, TRPA1 channels were most likely activated by elevated [Ca^2+^]_i_ or ROS. However, increased TRPA1 activity in ischaemia may also involve a change in Ca^2+^-mediated channel inhibition, or a shift in voltage-dependent gating to physiologically relevant values (half-maximal channel activation normally lies between +90 and +170 mV, but is shifted as low as -36 mV by agonists such as [Ca^2+^]_i_ or ROS).^51^ Accordingly, reducing VP_MAR_ duration, blocking TRPA1 channels, or limiting diastolic [Ca^2+^]_i_ and ROS would reduce the incidence of systolic MAR. Additionally, K_ATP_ channels are also mechano-sensitive. While quiescent under physiological conditions, if activated through a reduction in ATP levels during ischaemia or pharmacologically by PIN, the increase in their open probability with stretch is potentiated.^52^ The additional stretch-induced change in repolarisation (while potentially small, due to the transient nature of systolic stretch) may also be pro-arrhythmic, by further shortening APD and refractoriness, and thereby increasing the period of excitability between normal excitations.

The finding that dyscoupling of RRC, increased TRPA1 channel activity, and elevated [Ca^2+^]_i_ or ROS can lead to systolic MAR may have important implications for anti-arrhythmic treatments in various cardiac pathologies where these changes occur.^38^ In the case of myocardial ischaemia, experimental^8,53^ and computational^54^ studies have suggested that ventricular arrhythmias originating at the ischaemic border are in part driven by systolic stretch-induced excitation and altered repolarisation, caused by mechano-sensitive ion channels interacting with ischaemic changes in the electrophysiological substrate. However, the molecular mechanisms responsible for systolic MAR have not been identified. As it is known that during ischaemia: (i) dyscoupling of RRC occurs;^8,9^ (ii) TRPA1 channels are activated;^44,55^ and (iii) [Ca^2+^]_i_ and ROS levels are increased,^23^ these various factors may represent the mechanisms underlying systolic MAR in ischaemic myocardium.

Previous efforts to elucidate the presence and mechanisms of systolic MAR in the intact heart have been limited by the spatio-temporal complexity of cardiac electro-mechanical activity. In fact, other cases of MAR deemed to be systolic, such as *Commotio cordis* (which can occur in healthy tissue),^56^ may instead depend on stretch-induced excitation initiating from CM that have already returned to electrical diastole (so, at the CM level, this response may be driven by diastolic rather than systolic events, even if that may not be intuitively evident from gross electrophysiological observations in the intact heart, such as the electrocardiogram).^21^ However, in cardiac pathologies involving RRC dyscoupling, systolic stretch may underlie focal arrhythmogenesis, such as in failing human hearts with β-adrenergic alterations that result in an increase in the VP_MAR_.^57^ Additionally, even in situations that may be driven primarily by diastolic stretch (such as *Commotio cordis*), K_ATP_ channels may be a potential target to decrease MAR severity.^58^

In conclusion, as the molecular mechanisms responsible for stretch-induced changes in cardiac electrical activity have remained elusive,^5^ the identification of RRC dyscoupling and TRPA1 channels as a potential contributors in diseases in which their occurrence, expression, or activity is increased is an exciting development. Overall, the findings from this study suggest that dyscoupling of RRC in general, and TRPA1 channel activity in particular, may represent novel therapeutic targets for the prevention of systolic MAR in cardiac disease.

## STATEMENTS

## NON-STANDARD ABBREVIATIONS AND ACRONYMS

AITC: allyl isothiocyanate
CaTD: Ca^2+^ transient duration
APD: AP duration
CF: carbon fibre
CM: cardiomyocytes
CTRL: control
DPI: diphenyleneiodonium
ECC: excitation-contraction coupling
K_ATP_: ATP-sensitive potassium
MAR: mechano-arrhythmogenesis
MEC: mechano-electric coupling
NAC: N-acetyl-L-cysteine
PIN: pinacidil
RRC: repolarisation-relaxation coupling
SI: simulated ischaemia
TRPA1: transient receptor potential kinase ankyrin 1
T_RRC_: duration of RRC
V_m_: membrane potential
VP_MAR_: vulnerable period for MAR

## Acknowledgements.

The authors thank Gentaro Iribe (Asahikawa Medical University, Asahikawa, Japan) and Keiko Kaihara (Okayama University, Okayama, Japan) for technical assistance with cell stretch, and Ilija Uzelac (Georgia Institute of Technology, Atlanta, GA) for providing the control box for ratiometric imaging. CF were a gift from Jean-Yves LeGuennec (deceased).

